# Identification of the putative gene encoding the RNA subunit of telomerase in *Malassezia* clade species through comparative genomic analysis

**DOI:** 10.1101/2024.09.26.615164

**Authors:** Juan Antonio Sanpedro Luna, Patricia Sánchez Alonso

## Abstract

Maintaining telomere length and genomic stability in eukaryotic cells is dependent on telomerase, a ribonucleoprotein with reverse transcriptase activity that counteracts the end replication problem. Its core components include TERT (telomerase reverse transcriptase) and TER (telomerase RNA). This study focused on the identification of the putative *TER1* gene that encodes the telomerase RNA subunit in 19 *Malassezia* species via comparative genomics. Unlike the corresponding genes in other yeasts and filamentous fungi, this gene is compact and uniform in length, resembling similar genes in vertebrates, while retaining characteristic elements that are crucial for TER and snRNA biogenesis and activity. These elements include the template domain, pseudoknot domain, CR4/5 domain, and a putative Sm-binding site. Furthermore, transcriptomic analysis confirmed the transcriptional activity of the putative *TER1* gene, reinforcing its potential functional significance within the *Malassezia* clade. These results establish a robust basis for future experimental investigations to verify the functionality and regulatory mechanisms of this gene, particularly its role in telomere maintenance and broader cellular processes.

## Introduction

Telomerase is a ribonucleoprotein with reverse transcriptase activity that catalyzes the *de novo* synthesis of telomeric repeats at the linear ends of eukaryotic chromosomes [1]. Telomerase activity addresses the end replication problem [2, 3] by elongating telomeres. The maintenance of telomere length by telomerase allows telomeres to act as protective caps on chromosome ends and prevent DNA repair machinery from recognizing these ends as double-strand breaks. Therefore, the activity of telomerase is critical and contributes to the maintenance of genomic stability by preventing the erosion of chromosomal end sequences and fusion-bridge-breakage cycles [4, 5]. The core components that form the catalytic core of the complex required for the restoration of catalytic activity *in vitro* are the protein subunit TERT (<u>Te</u>lomerase <u>R</u>everse <u>T</u>ranscriptase), which functions as a reverse transcriptase, and the subunit TER (<u>Te</u>lomerase <u>R</u>NA), which serves as a template for reverse transcription and a scaffolding site for components involved in complex activity [6].

In addition to its telomeric function, TER may have extratelomeric functions in different organisms, including the regulation of key biological processes. It has been reported that this subunit plays an antiapoptotic role in response to cellular stress [7, 8], participates in the DNA damage response [9, 10], and regulates gene expression and cell differentiation [11, 12]; moreover, in dimorphic fungi, TER has been shown to participate in regulating genes that are involved in the life cycle and pathogenic development [13, 14]. These findings show that the telomerase RNA subunit plays broader roles in cellular biology beyond its function in telomere stability.

The identification of genes related to telomeres and telomerase in various organisms has been an area of interest in biological research. Recent progress has been made in understanding the genomic structure of the genus *Malassezia*. These advances provide valuable information about the organization of chromosomal ends and the components involved in their maintenance [15]. Moreover, the study of telomerase components and their functions has yet to be explored. The genus *Malassezia* is classified within the division Basidiomycota, belonging to the class Malasseziomycetes [16]; this genus is composed of a group of dimorphic yeast dependent on lipids [17], which inhabit the skin surface of humans and a wide range of warm-blooded animals [18] and constitute the predominant component of the cutaneous mycobiome in humans [19]. Species in this clade typically have a commensal lifestyle, although they are also involved in the development of dermatological diseases [20], and their possible mutualistic mechanism has been described [21]. Despite the difficulties involved in growing *Malassezia* strains under laboratory conditions, their presence has been detected, via molecular methods, in diverse habitats, such as deep-sea ecosystems [22], Antarctic soils [23], corals, sponges, and cone snails [24]. This highlights the great ecological diversity and adaptability of *Malassezia*.

Unlike other phylogenetically related and extensively studied fungi, such as Ustilaginomycetes [16], this clade contains significantly smaller genomes ranging from 7— 9 Mb, with genes distributed across a variable number of chromosomes ranging from six to nine [25, 26]. The recent availability of genomic sequences from various *Malassezia* species has provided a valuable opportunity to more comprehensively explore telomere biogenesis in this group of fungi, while the differences in life cycle, organization, and genetic content compared with those of Ustilaginomycetes provide a unique opportunity to study the function and evolution of the components involved in telomere maintenance in these organisms.

## Materials and Methods

### Identification of the putative *TER1* gene in *Malassezia* species

The identification of the putative *TER1* gene was performed following a previously described method [27]. A comprehensive search for all regions containing at least one and a half copies of the expected template sequence (5’-CACTAA-3’) in any possible permutation within the genome of *M. globosa* (GenBank accession: GCA_010232095.1) was conducted. The candidate loci, along with short flanking sequences on each side, were subsequently used as search queries to identify orthologous sequences in the genomes of 18 additional *Malassezia* species (S1 Table) available to date via BLASTn [28]. Next, multiple sequence alignments were performed to identify conserved motifs within the putative *TER1* gene. The sequences obtained from the orthologous genes of the different *Malassezia* species were aligned via MAFFT [29] with the G-INS-i strategy, and the --unalignlevel 0.8 and --leavegappyregion options. The alignment was visualized with BioEdit software [30], allowing for a comprehensive analysis of the conserved regions.

### Secondary Structure Prediction

To determine the secondary structure, multiple sequence alignments were utilized to identify conserved regions extracted from the *M. globosa* sequence. The total length of the transcript was estimated on the basis of the first and last well-conserved motifs within the intergenic region. This region was extracted and analyzed with Mfold [31] to predict its secondary structure. The obtained structure was subsequently supported by the consensus secondary structures obtained via RNAalifold [32] from the aligned sequence set, and the conservation of the regions and their positions within the predicted structure was subsequently assessed via alignment analysis.

### Transcriptomic profiling of the putative *TER1* gene

To obtain insights into the transcriptional activity of the putative *TER1* gene, RNA-seq data available for *M. globosa* were downloaded (accession number: SRR2072129-SRR2072131), and the quality of the raw paired reads was assessed via FastQC v0.11.9 [33]. The removal of ribosomal RNA reads was subsequently performed with SortMeRNA software v4.2.0 [34]. The trimming and removal of low-quality reads were performed with Trimmomatic v0.39 [35], and reads with scores greater than 30 and a minimum length of 25 nt were retained for further analysis. Next, *de novo* assembly was performed with Trinity v2.15.1 [36] on the processed reads with the following parameters: --min_kmer_cov=2 -- min_contig_length=100 --jaccard_clip --SS_lib_type FR. The reconstructed transcripts and processed reads were mapped to the reference genome with GMAP v23-02-17 [37] and Bowtie 2 v2.4.4 [38], respectively. The alignments were visualized and explored with IGV (Integrative Genomics Viewer) software v2.16.0 [39], enabling a comprehensive examination of the putative gene and its genomic surroundings.

### Phylogenetic tree construction

A phylogenetic tree was constructed using the putative *TER1* gene sequences from the 19 *Malassezia* species. The sequences were analyzed with MEGA11 software v11.0.13 [40], employing the maximum likelihood approach with the HKY nucleotide substitution model. To assess the robustness of the tree groupings, 500 bootstrap replicates were performed.

## Results

### Identification of the putative *TER1* gene

The mining of template domain sequences within the *M. globosa* genome and the search for orthologous regions allowed the identification of a unique intergenic region that was moderately conserved in all the examined *Malassezia* species. In all the examined species, such as *M. globosa*, this region was found to be flanked at its 3’ end by a reading frame encoding a protein of unknown function that contained the conserved DUF1115 domain. Although the presence of the *NDK1* gene, which encodes nucleoside diphosphate kinase, was observed at the 5’ end, in at least 4 species, a reading frame was identified interspersed between *NDK1* and the TER loci, which encodes a cysteine hydrolase (S1 Fig). Once the orthologous regions of the TER locus from *M. globosa* were identified, the sequences were aligned and examined to identify conserved motifs. Initially, the conserved motif of the putative template domain was examined, as shown in Fig 1, and in some cases, point mutations were identified. These mutations, however, were consistent with the expected template sequence for the telomeric repeat previously reported for each species [15] (Fig 2).

**Fig 1.**
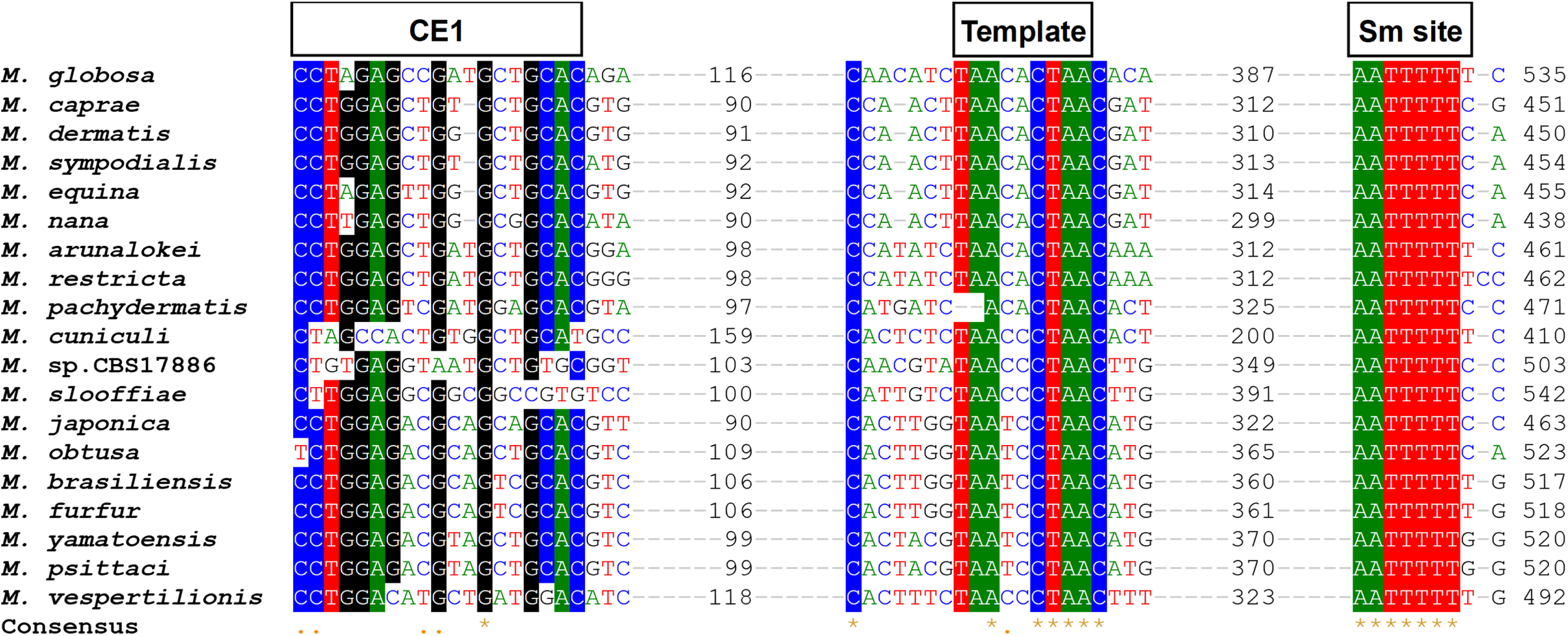
Multiple sequence alignment of the putative *TER1* gene in *Malassezia* clade species. In the multiple sequence alignment of the putative *TER1* gene in 19 *Malassezia* species, in addition to the conserved template domain, upstream, a conserved element (CE1) of unknown function was identified. Toward the 3’ end of the intergenic region, a well-conserved motif resembling a Sm-binding site was identified. These regions were considered the start and end of the transcript. Conserved regions present in at least 75% of the species are highlighted with a colored background. Structural domains were identified via boxed annotations.

**Fig 2.**
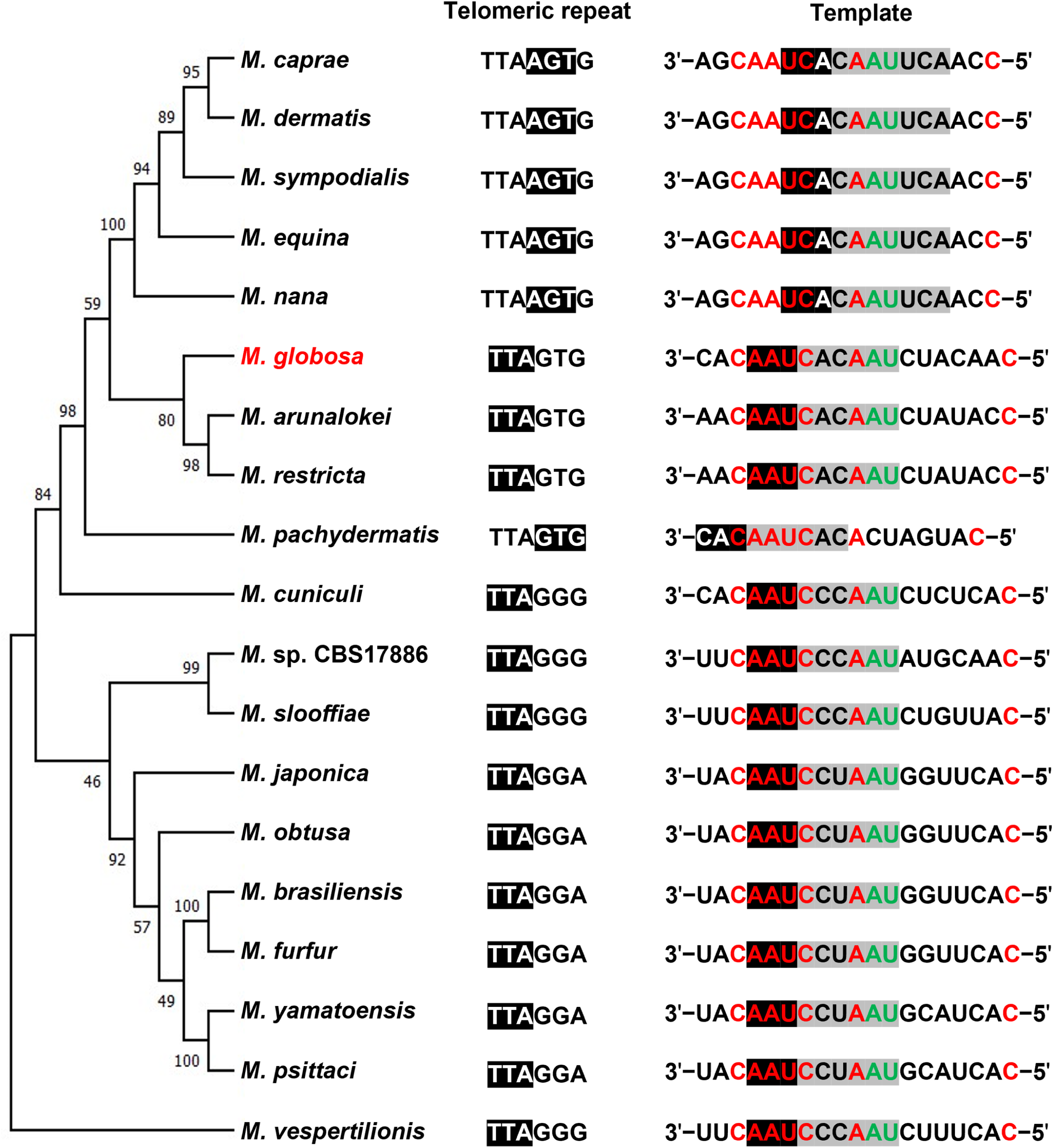
Telomeric repeats and phylogenetic tree of the putative *TER1* gene in *Malassezia* clade species. Evolutionary relationships of the *TER1* gene among *Malassezia* species, telomeric repeats, and sequences containing the template domain (Columns 1 and 2, respectively) are shown. Nucleotides conserved in all species are shown in red, while nucleotides conserved in all species except *M. pachydermatis* are shown in green. The alignment region is highlighted in black, and the template region for retrotranscription is highlighted in gray. Point mutations within the region cause sliding of the template sequence and the generation of a new telomeric repeat.

Toward the 5’ end of this domain, the first conserved region (Conserved element 1), located at an average length of 112 nt from the template domain, was identified. To determine its potential function, an analysis was conducted to search for promoter regions or regulatory sequences within this conserved motif. However, no regions associated with the regulation of the putative gene could be identified. Instead, it was considered that this conserved region might be part of a structural element of the mature transcript, Therefore, it was included in the sequence folding analysis. Toward the 3’ end, the consensus sequence 5’-AAUUUUU-3’, which is identical to the putative Sm-binding site of Ustilaginales [13, 14], was identified. Similarly, no conserved motifs suggesting processing by splicing were identified downstream of this domain in this clade. Therefore, these sequences were considered the start and end of the gene, respectively, resulting in an average length of 483 nt.

### The putative *TER1* gene contains essential elements conserved for its biogenesis

Once the estimated length of the transcript was established, the folding of the sequence was analyzed with Mfold, resulting in a structural model of TER from *M. globosa* that resembles the structure of vertebrate and fungal TER [41, 42]. This model contains structural elements required for subunit activity, which, according to the multiple sequence alignment, constitute conserved motifs that are essential for the biogenesis and activity of the mature transcript (S2 Fig). In addition to the template domain and the putative Sm-binding site [43–45], other conserved domains were identified, such as the pseudoknot and the CR4/5 domain near the 3’ end that contains a structure analogous to the P6.1 helix (Fig 3) [42, 46].

**Fig 3.**
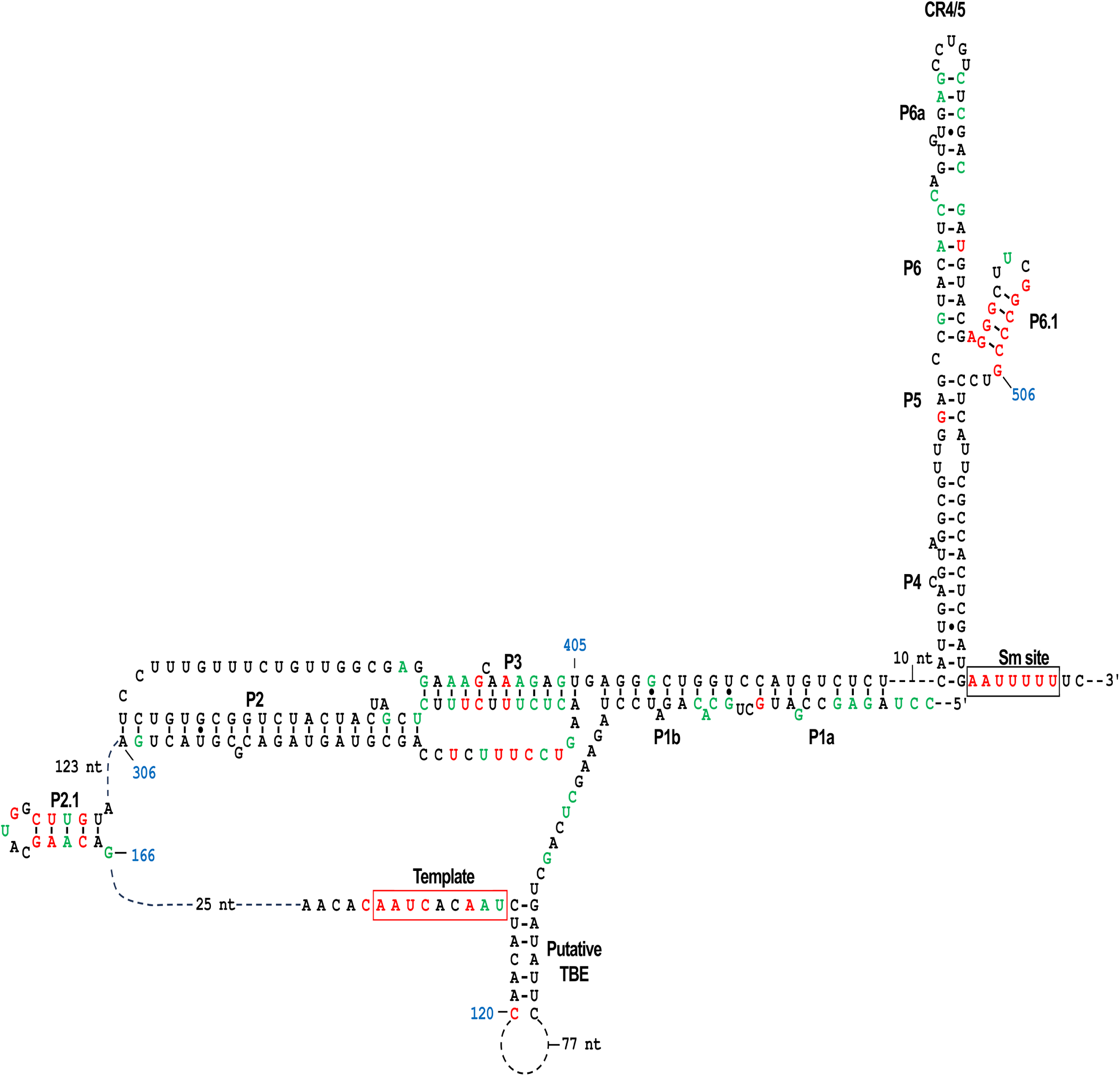
Secondary structure of the telomerase RNA subunit in *M. globosa*. The mature TER subunit in *M. globosa* has an approximate length of 535 nt and can fold into a secondary structure that contains the structural domains required for subunit activity, including the template domain, TBE, pseudoknot, CR4/5 domain, and Sm-binding site. The conserved nucleotides in all the examined species are depicted in red. Nucleotides conserved in at least 75% of the species are depicted in green.

The template domain was located in a single-stranded region, and adjacent to this element, the putative TBE structure was observed to be composed of a poorly conserved sequence. Consistent with the requirement for the formation of a triple helix in the pseudoknot domain, which is typically stabilized by three consecutive segments of U-A·U triple bases [47, 48], a moderately conserved region with similar characteristics was identified downstream of the template domain. This region has the potential to transcribe sequences with high contents of U and A residues. This region forms a stem that corresponds to stem 2 of the pseudoknot domain, as it is formed by U-A base pairs interspersed with C-G pairings. In addition, conserved residues of U and C are present in loop 1, suggesting the potential to form U-A·U triple bases. Additionally, between the T/PK domains, similar to those of basal metazoans [46], a conserved stem called P2.1 was identified. Along with the structures of P3 and P6.1, P2.1 was located in the consensus secondary structure obtained with RNAalifold.

### Transcriptional activity of the putative *TER1* gene

To obtain information about the transcriptional activity of the intergenic region where the putative *TER1* gene was located and its potential length, paired-end reads were mapped to the reference genome and used for *de novo* assembly. Following the processing of raw reads, a total of 47,746,223 paired-end reads were retained and utilized for assembly, resulting in a set of 21,968 unique transcripts. The average length of these transcripts was 893 nucleotides. Detailed assembly statistics are provided in S2 Table. Additionally, comprehensive results of raw data processing and sequence alignment statistics against the *de novo* transcriptome assembly are presented in S3 Table. With respect to the transcription of the putative TER locus, a single transcript containing the complete sequence of the putative gene was identified in the assembled transcripts. Additionally, the presence of aligned reads throughout the intergenic region was observed, with a notable increase in the region corresponding to the sequence containing the structural putative domains of TER. Interestingly, this transcript is 1,756 bp long, and its transcription start site is located upstream of the flanking 5’ end of the gene (Fig 4). The gene consists of 2 exons and corresponds to the ortholog of *NDK1*, which encodes a well-characterized nucleoside diphosphate kinase. These results suggest that this putative gene performs a biological function by revealing the presence of transcribed sequences in the intergenic region where the putative *TER1* gene is located that cover all the essential elements for activity (Fig 4).

**Fig 4.**
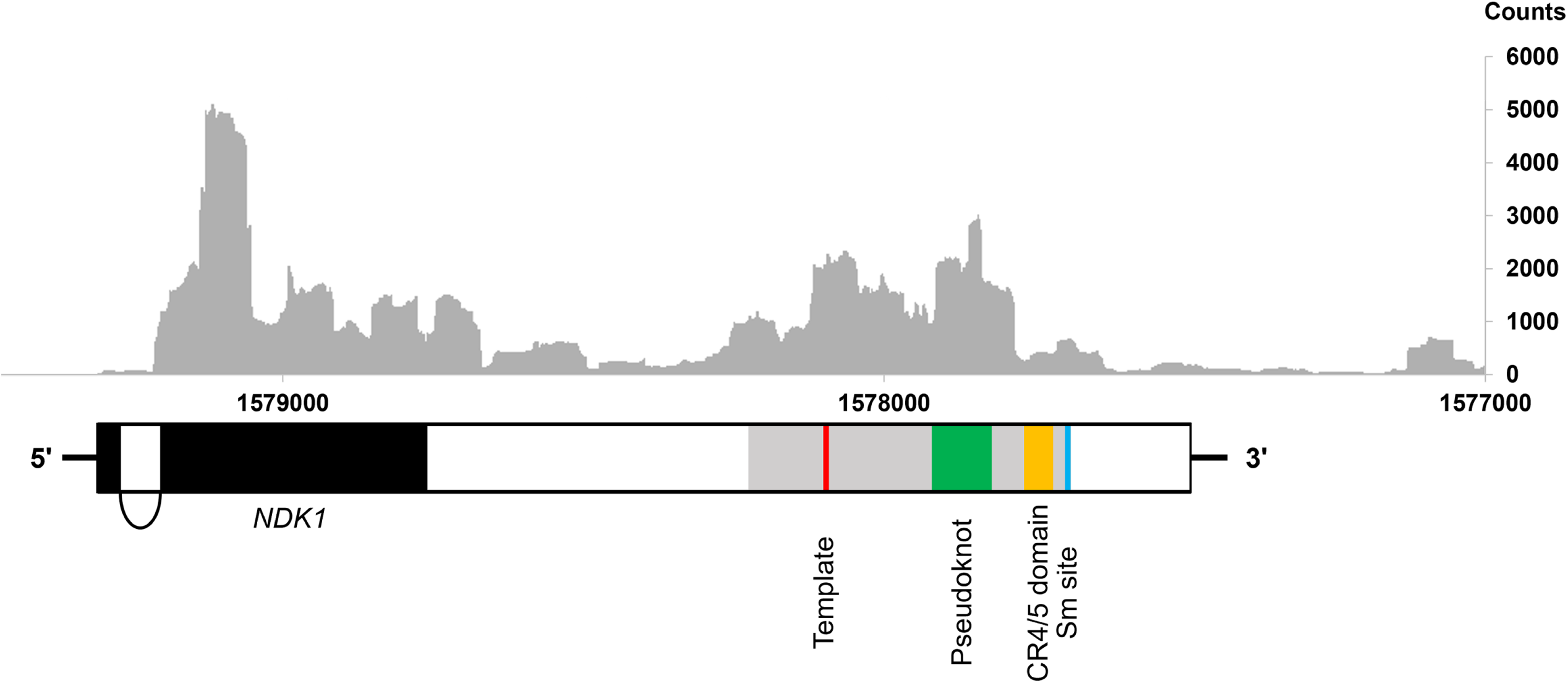
Representation of the putative *TER1* gene identified on chromosome 1 of *M. globosa*. The locations of the structural domains within the putative *TER1* gene (gray rectangle) are represented by colored boxes. The upper part of the figure shows the density distribution of RNA-seq reads along the gene. The results of the RNA-seq read distribution confirmed the transcriptional capability of the intergenic region containing the putative *TER1* gene, suggesting its potential to generate functional RNA in *M. globosa*.

## Discussion

The availability of public databases and the increasing number of genomic sequences from Basidiomycota provide a valuable opportunity to continue studying TER in a phylogenetically diverse and understudied group of fungi. In this case, the bioinformatics approach allowed for the successful identification of the putative *TER1* gene within the *Malassezia* clade. The comparative study of the structure and sequence of TER between Basidiomycota and Ascomycota will help identify possible adaptations and evolutionary changes that have occurred in telomerase over time and in different fungal lineages.

While the identification of TER through conventional alignments of the gene encoding the catalytic subunit followed by precipitation has proven to be an effective technique [45, 49], this approach is complex, highlighting that, despite its success, it has inherent challenges and limitations. On the other hand, although the computational approach simplifies identification, it is crucial to recognize that this strategy still has its own complexities and limitations. This approach is restricted to individual searches within narrow phylogenetic groups owing to the inherent plasticity of noncoding transcripts and the rapid evolution of genomes, as evidenced by the diversity of sequences that hinder reliable alignments and their identification in Saccharomycetes [50].

Interestingly, the putative *TER1* gene identified in this clade is similar in length to the TER gene in vertebrates [41], but unlike the latter, the presence of the putative Sm-binding site in this gene indicates that 3’ end processing follows the typical snRNA pathway of fungi and yeasts [43, 51]. On the other hand, the length of this gene is different from that of *ter1* in Ustilaginales, which has a more heterogeneous sequence and a greater average length [13, 14]; this could be explained by the occurrence of more events of sequence accumulation between essential motifs, as well as transcriptional fusion with the locus UMAG_03168, which encodes a protein involved in meiosis [13], whose homolog in *M. globosa* is located at a distal position on another chromosome (S3 Fig). In addition, the reconstruction of a continuous transcript containing a coding gene and TER structural elements, along with a divergent intergenic sequence at the 5’ end and the absence of conserved motifs upstream of the TER loci, suggests that, as in the Ustilaginales, the mature TER transcript in the *Malassezia* clade also originates from a coding gene. Although limited information on TER in Basidiomycota is available to date, the conservation of a more compact gene linked to the ancestral form of TER suggests that TER-intron fusion at the 3’ end in the early Ascomycota lineages and the 3’ end processing mediated by the spliceosome [52] is exclusive to the phylum Ascomycota and occurred after the divergence from Basidiomycota. Despite the conserved gene length, a notable change in the template domain sequence within this clade gives rise to four distinct telomeric repeats [15], which can be generated by alternate changes in the template domain, as observed in *M. pachydermatis*.

The formation of a new telomeric repeat can be directly attributed to point mutations observed in the template domain or to alignment slippage caused by mutations in the region surrounding the template domain. Furthermore, as shown by the phylogenetic analysis of the putative gene, the origin of these telomeric repeats is consistent with a common ancestor bearing the TTAGGG sequence (Fig 2). This, in turn, is consistent with the idea that this sequence possibly corresponds to the most ancestral template sequence of TER across diverse lineages [53, 54]. Simultaneously, the sustained utilization of templates exhibiting minor variations suggests the presence of discernible selective pressure, likely stemming from the imperative requirement for interaction with proteins associated with telomeres.

## Conclusions

In this study, we successfully identified the putative gene encoding the RNA subunit of telomerase in 19 *Malassezia* species. Similar to related genes in other yeasts and filamentous fungi, the putative Sm-binding site located at the 3’ end of this gene suggests its processing through the snRNA pathway. However, unlike similar genes in these other organisms, this gene is more compact and homogeneous in length, with an estimated size similar to those of related genes in vertebrates. Additionally, it contains the necessary structural elements that support its activity. The biological function of this gene was revealed by the analysis of RNA-seq data, confirming the transcriptional activity of the putative gene.

The implementation of bioinformatic analyses represents a highly efficient and agile alternative for obtaining biological insights and generating approaches that significantly simplify processes and guide the development of experimental strategies. The findings presented here provide a solid foundation for future research focused on experimentally confirming the function and activity of this gene in the context of telomere maintenance regulation and its potential involvement in other cellular processes. Ultimately, the study of telomerase regulation and evolution, which is based on information collected from diverse organisms, may contribute to a better understanding of the function and impact of telomerase components on cellular biology.

## Acknowledgments

The authors acknowledge Joseph Heitman, Márcia David-Palma and Marco Antonio Dias Coelho from the Department of Molecular Genetics and Microbiology, Duke University Medical Center, Durham, North Carolina, USA, for allowing the use of sequences for this analysis and for the useful comments to the work and to and Candelario Vazquez Cruz from the Benemérita Universidad Autónoma de Puebla, Puebla, Mexico for useful discussions of the work.

## Supporting information

**S1 Fig. Synteny map of *Malassezia* clade species around the putative *TER1* gene**. The region shows the conservation of genes flanking the *TER1* gene, highlighting the consistent arrangement of genetic elements. The reference genes were selected on the basis of the annotation of previously reported genomes [15]. Gene models were reviewed, and manual corrections were applied as necessary for all species.

**S2 Fig. Multiple sequence alignment of 19 TERs from *Malassezia* clade species.** Multiple sequence alignment of 19 TER sequences from species of the *Malassezia* clade was performed with MAFFT. The nucleotides conserved in ≥75% of the examined species are represented by a colored background.

**S3 Fig. Emi1 homolog identification.** Multiple alignments of isoforms A and B of Emi1 from *U. maydis* and the homologous protein were identified in the genome of *M. globosa* (chromosome 3, locus MGL_0937). The sequences were aligned using the ClustalW algorithm [55]. Conserved residues are highlighted with colored backgrounds.

S1 Table. Locations of the putative TER loci among the species of the *Malassezia* clade. S2 Table. Trinity *de novo* assembly statistics.

**S3 Table. Filtering of raw data and sequence mapping.**

## Data Availability

The genome and annotation data of *M. globosa* CBS 7966, utilized as a reference, were obtained from NCBI (GCA_010232095.1 and GCA_000181695.2, respectively). The RNA-seq data of *M. globosa* used for the analysis were retrieved from NCBI (accession numbers: SRR2072129-SRR2072131). The scripts and files utilized for both the assembly process and the corresponding statistical analyses can be accessed at https://github.com/Antonio-Sanpedro/Transcriptome_assembly. Detailed information on the reference sequences used for orthologous identification and the putative *TER1* gene sequences deposited as third-party annotation (TPA) entries in GenBank (PENDING) can be found in S1 Table.

## References

1. Greider CW, Blackburn EH. A telomeric sequence in the RNA of Tetrahymena telomerase required for telomere repeat synthesis. Nature. 1989;337(6205):331-7. Epub 1989/01/26. doi: 10.1038/337331a0. PubMed PMID: 2463488.

2. Olovnikov AM. [The immune response and the process of marginotomy in lymphoid cells]. Vestn Akad Med Nauk SSSR. 1972;27(12):85–7. Epub 1972/01/01. PubMed PMID: 4204758.

3. Watson JD. Origin of concatemeric T7 DNA. Nat New Biol. 1972;239(94):197–201. Epub 1972/10/18. doi: 10.1038/newbio239197a0. PubMed PMID: 4507727.

4. Ducray C, Pommier JP, Martins L, Boussin FD, Sabatier L. Telomere dynamics, end-to-end fusions and telomerase activation during the human fibroblast immortalization process. Oncogene. 1999;18(29):4211–23. Epub 1999/08/06. doi: 10.1038/sj.onc.1202797. PubMed PMID: 10435634.

5. Hackett JA, Feldser DM, Greider CW. Telomere dysfunction increases mutation rate and genomic instability. Cell. 2001;106(3):275–86. Epub 2001/08/18. doi: 10.1016/s0092-8674(01)00457-3. PubMed PMID: 11509177.

6. Nguyen THD, Tam J, Wu RA, Greber BJ, Toso D, Nogales E, et al. Cryo-EM structure of substrate-bound human telomerase holoenzyme. Nature. 2018;557(7704):190-5. Epub 2018/04/27. doi: 10.1038/s41586-018-0062-x. PubMed PMID: 29695869; PubMed Central PMCID: PMCPMC6223129.

7. Gazzaniga FS, Blackburn EH. An antiapoptotic role for telomerase RNA in human immune cells independent of telomere integrity or telomerase enzymatic activity. Blood. 2014;124(25):3675–84. Epub 2014/10/17. doi: 10.1182/blood-2014-06-582254. PubMed PMID: 25320237; PubMed Central PMCID: PMCPMC4263978.

8. Rubtsova M, Naraykina Y, Vasilkova D, Meerson M, Zvereva M, Prassolov V, et al. Protein encoded in human telomerase RNA is involved in cell protective pathways. Nucleic Acids Res. 2018;46(17):8966–77. Epub 2018/08/14. doi: 10.1093/nar/gky705. PubMed PMID: 30102362; PubMed Central PMCID: PMCPMC6158713.

9. Kedde M, le Sage C, Duursma A, Zlotorynski E, van Leeuwen B, Nijkamp W, et al. Telomerase-independent regulation of ATR by human telomerase RNA. J Biol Chem. 2006;281(52):40503-14. Epub 2006/11/14. doi: 10.1074/jbc.M607676200. PubMed PMID: 17098743.

10. Ting NS, Pohorelic B, Yu Y, Lees-Miller SP, Beattie TL. The human telomerase RNA component, hTR, activates the DNA-dependent protein kinase to phosphorylate heterogeneous nuclear ribonucleoprotein A1. Nucleic Acids Res. 2009;37(18):6105–15. Epub 2009/08/07. doi: 10.1093/nar/gkp636. PubMed PMID: 19656952; PubMed Central PMCID: PMCPMC2764450.

11. Li S, Crothers J, Haqq CM, Blackburn EH. Cellular and gene expression responses involved in the rapid growth inhibition of human cancer cells by RNA interference-mediated depletion of telomerase RNA. J Biol Chem. 2005;280(25):23709–17. Epub 2005/04/16. doi: 10.1074/jbc.M502782200. PubMed PMID: 15831499.

12. Alcaraz-Perez F, Garcia-Castillo J, Garcia-Moreno D, Lopez-Munoz A, Anchelin M, Angosto D, et al. A non-canonical function of telomerase RNA in the regulation of developmental myelopoiesis in zebrafish. Nat Commun. 2014;5:3228. Epub 2014/02/06. doi: 10.1038/ncomms4228. PubMed PMID: 24496182.

13. Logeswaran D, Li Y, Akhter K, Podlevsky JD, Olson TL, Forsberg K, et al. Biogenesis of telomerase RNA from a protein-coding mRNA precursor. Proc Natl Acad Sci U S A. 2022;119(41):e2204636119. Epub 2022/10/06. doi: 10.1073/pnas.2204636119. PubMed PMID: 36197996; PubMed Central PMCID: PMCPMC9564094.

14. Sanpedro-Luna JA, Jacinto-Vazquez JJ, Anastacio-Marcelino E, Posadas-Gutierrez CM, Olmos-Pineda I, Gonzalez-Bernal JA, et al. Telomerase RNA plays a major role in the completion of the life cycle in Ustilago maydis and shares conserved domains with other Ustilaginales. PLoS One. 2023;18(3):e0281251. Epub 2023/03/24. doi: 10.1371/journal.pone.0281251. PubMed PMID: 36952474; PubMed Central PMCID: PMCPMC10035886.

15. Coelho MA, Ianiri G, David-Palma M, Theelen B, Goyal R, Narayanan A, et al. Frequent transitions in mating-type locus chromosomal organization in Malassezia and early steps in sexual reproduction. Proc Natl Acad Sci U S A. 2023;120(32):e2305094120. Epub 2023/07/31. doi: 10.1073/pnas.2305094120. PubMed PMID: 37523560; PubMed Central PMCID: PMCPMC10410736.

16. Wang QM, Theelen B, Groenewald M, Bai FY, Boekhout T. Moniliellomycetes and Malasseziomycetes, two new classes in Ustilaginomycotina. Persoonia. 2014;33:41–7. Epub 2015/03/05. doi: 10.3767/003158514X682313. PubMed PMID: 25737592; PubMed Central PMCID: PMCPMC4312936.

17. Wu G, Zhao H, Li C, Rajapakse MP, Wong WC, Xu J, et al. Genus-Wide Comparative Genomics of Malassezia Delineates Its Phylogeny, Physiology, and Niche Adaptation on Human Skin. PLoS Genet. 2015;11(11):e1005614. Epub 2015/11/06. doi: 10.1371/journal.pgen.1005614. PubMed PMID: 26539826; PubMed Central PMCID: PMCPMC4634964 shampoo.

18. Gueho E, Midgley G, Guillot J. The genus Malassezia with description of four new species. Antonie Van Leeuwenhoek. 1996;69(4):337–55. Epub 1996/05/01. doi: 10.1007/BF00399623. PubMed PMID: 8836432.

19. Findley K, Oh J, Yang J, Conlan S, Deming C, Meyer JA, et al. Topographic diversity of fungal and bacterial communities in human skin. Nature. 2013;498(7454):367-70. Epub 2013/05/24. doi: 10.1038/nature12171. PubMed PMID: 23698366; PubMed Central PMCID: PMCPMC3711185.

20. DeAngelis YM, Gemmer CM, Kaczvinsky JR, Kenneally DC, Schwartz JR, Dawson TL, Jr. Three etiologic facets of dandruff and seborrheic dermatitis: Malassezia fungi, sebaceous lipids, and individual sensitivity. J Investig Dermatol Symp Proc. 2005;10(3):295–7. Epub 2005/12/31. doi: 10.1111/j.1087-0024.2005.10119.x. PubMed PMID: 16382685.

21. Li H, Goh BN, Teh WK, Jiang Z, Goh JPZ, Goh A, et al. Skin Commensal Malassezia globosa Secreted Protease Attenuates Staphylococcus aureus Biofilm Formation. J Invest Dermatol. 2018;138(5):1137–45. Epub 2017/12/17. doi: 10.1016/j.jid.2017.11.034. PubMed PMID: 29246799.

22. Le Calvez T, Burgaud G, Mahe S, Barbier G, Vandenkoornhuyse P. Fungal diversity in deep- sea hydrothermal ecosystems. Appl Environ Microbiol. 2009;75(20):6415–21. Epub 2009/07/28. doi: 10.1128/AEM.00653-09. PubMed PMID: 19633124; PubMed Central PMCID: PMCPMC2765129.

23. Fell JW, Scorzetti G, Connell L, Craig SJSB, Biochemistry. Biodiversity of micro-eukaryotes in Antarctic Dry Valley soils with < 5% soil moisture. 2006;38(10):3107–19.

24. Amend A. From dandruff to deep-sea vents: Malassezia-like fungi are ecologically hyper- diverse. PLoS Pathog. 2014;10(8):e1004277. Epub 2014/08/22. doi: 10.1371/journal.ppat.1004277. PubMed PMID: 25144294; PubMed Central PMCID: PMCPMC4140847.

25. Boekhout T, Kamp M, Gueho E. Molecular typing of Malassezia species with PFGE and RAPD. Med Mycol. 1998;36(6):365–72. Epub 1999/04/17. doi: 10.1080/02681219880000581. PubMed PMID: 10206745.

26. Sankaranarayanan SR, Ianiri G, Coelho MA, Reza MH, Thimmappa BC, Ganguly P, et al. Loss of centromere function drives karyotype evolution in closely related Malassezia species. Elife. 2020;9. Epub 2020/01/21. doi: 10.7554/eLife.53944. PubMed PMID: 31958060; PubMed Central PMCID: PMCPMC7025860.

27. Chakrabarti K, Pearson M, Grate L, Sterne-Weiler T, Deans J, Donohue JP, et al. Structural RNAs of known and unknown function identified in malaria parasites by comparative genomics and RNA analysis. RNA. 2007;13(11):1923–39. Epub 2007/09/29. doi: 10.1261/rna.751807. PubMed PMID: 17901154; PubMed Central PMCID: PMCPMC2040097.

28. Altschul SF, Gish W, Miller W, Myers EW, Lipman DJ. Basic local alignment search tool. J Mol Biol. 1990;215(3):403–10. doi: 10.1016/S0022-2836(05)80360-2. PubMed PMID: 2231712.

29. Katoh K, Misawa K, Kuma K, Miyata T. MAFFT: a novel method for rapid multiple sequence alignment based on fast Fourier transform. Nucleic Acids Res. 2002;30(14):3059–66. Epub 2002/07/24. doi: 10.1093/nar/gkf436. PubMed PMID: 12136088; PubMed Central PMCID: PMCPMC135756.

30. Hall T A. BioEdit : a user-friendly biological sequence alignment editor and analysis program for Windows 95/98/NT. Nucleic Acids Symp Ser. 1999;41:95–8.

31. Zuker M. Mfold web server for nucleic acid folding and hybridization prediction. Nucleic Acids Res. 2003;31(13):3406–15. Epub 2003/06/26. doi: 10.1093/nar/gkg595. PubMed PMID: 12824337; PubMed Central PMCID: PMCPMC169194.

32. Bernhart SH, Hofacker IL, Will S, Gruber AR, Stadler PF. RNAalifold: improved consensus structure prediction for RNA alignments. BMC Bioinformatics. 2008;9:474. Epub 2008/11/19. doi: 10.1186/1471-2105-9-474. PubMed PMID: 19014431; PubMed Central PMCID: PMCPMC2621365.

33. Andrews S., Fast QC: A quality control tool for high throughput sequence data. Available online at:. 2010;https://www.bioinformatics.babraham.ac.uk/projects/fastqc/.

34. Kopylova E, Noe L, Touzet H. SortMeRNA: fast and accurate filtering of ribosomal RNAs in metatranscriptomic data. Bioinformatics. 2012;28(24):3211–7. Epub 2012/10/17. doi: 10.1093/bioinformatics/bts611. PubMed PMID: 23071270.

35. Bolger AM, Lohse M, Usadel B. Trimmomatic: a flexible trimmer for Illumina sequence data. Bioinformatics. 2014;30(15):2114–20. doi: 10.1093/bioinformatics/btu170. PubMed PMID: 24695404; PubMed Central PMCID: PMCPMC4103590.

36. Grabherr MG, Haas BJ, Yassour M, Levin JZ, Thompson DA, Amit I, et al. Full-length transcriptome assembly from RNA-Seq data without a reference genome. Nat Biotechnol. 2011;29(7):644–52. Epub 2011/05/17. doi: 10.1038/nbt.1883. PubMed PMID: 21572440; PubMed Central PMCID: PMCPMC3571712.

37. Wu TD, Watanabe CK. GMAP: a genomic mapping and alignment program for mRNA and EST sequences. Bioinformatics. 2005;21(9):1859–75. Epub 2005/02/25. doi: 10.1093/bioinformatics/bti310. PubMed PMID: 15728110.

38. Langmead B, Salzberg SL. Fast gapped-read alignment with Bowtie 2. Nat Methods. 2012;9(4):357–9. Epub 2012/03/06. doi: 10.1038/nmeth.1923. PubMed PMID: 22388286; PubMed Central PMCID: PMCPMC3322381.

39. Robinson JT, Thorvaldsdottir H, Winckler W, Guttman M, Lander ES, Getz G, et al. Integrative genomics viewer. Nat Biotechnol. 2011;29(1):24-6. Epub 2011/01/12. doi: 10.1038/nbt.1754. PubMed PMID: 21221095; PubMed Central PMCID: PMCPMC3346182.

40. Tamura K, Stecher G, Kumar S. MEGA11: Molecular Evolutionary Genetics Analysis Version 11. Mol Biol Evol. 2021;38(7):3022–7. Epub 2021/04/24. doi: 10.1093/molbev/msab120. PubMed PMID: 33892491; PubMed Central PMCID: PMCPMC8233496.

41. Chen JL, Blasco MA, Greider CW. Secondary structure of vertebrate telomerase RNA. Cell. 2000;100(5):503–14. Epub 2000/03/18. doi: 10.1016/s0092-8674(00)80687-x. PubMed PMID: 10721988.

42. Qi X, Li Y, Honda S, Hoffmann S, Marz M, Mosig A, et al. The common ancestral core of vertebrate and fungal telomerase RNAs. Nucleic Acids Res. 2013;41(1):450–62. Epub 2012/10/25. doi: 10.1093/nar/gks980. PubMed PMID: 23093598; PubMed Central PMCID: PMCPMC3592445.

43. Seto AG, Zaug AJ, Sobel SG, Wolin SL, Cech TR. Saccharomyces cerevisiae telomerase is an Sm small nuclear ribonucleoprotein particle. Nature. 1999;401(6749):177-80. Epub 1999/09/18. doi: 10.1038/43694. PubMed PMID: 10490028.

44. Box JA, Bunch JT, Tang W, Baumann P. Spliceosomal cleavage generates the 3’ end of telomerase RNA. Nature. 2008;456(7224):910-4. Epub 2008/12/05. doi: 10.1038/nature07584. PubMed PMID: 19052544.

45. Leonardi J, Box JA, Bunch JT, Baumann P. TER1, the RNA subunit of fission yeast telomerase. Nat Struct Mol Biol. 2008;15(1):26–33. Epub 2007/12/25. doi: 10.1038/nsmb1343. PubMed PMID: 18157152.

46. Logeswaran D, Li Y, Podlevsky JD, Chen JJ. Monophyletic Origin and Divergent Evolution of Animal Telomerase RNA. Mol Biol Evol. 2021;38(1):215–28. Epub 2020/08/10. doi: 10.1093/molbev/msaa203. PubMed PMID: 32770221; PubMed Central PMCID: PMCPMC8480181.

47. Theimer CA, Blois CA, Feigon J. Structure of the human telomerase RNA pseudoknot reveals conserved tertiary interactions essential for function. Mol Cell. 2005;17(5):671–82. Epub 2005/03/08. doi: 10.1016/j.molcel.2005.01.017. PubMed PMID: 15749017.

48. Ulyanov NB, Shefer K, James TL, Tzfati Y. Pseudoknot structures with conserved base triples in telomerase RNAs of ciliates. Nucleic Acids Res. 2007;35(18):6150–60. Epub 2007/09/11. doi: 10.1093/nar/gkm660. PubMed PMID: 17827211; PubMed Central PMCID: PMCPMC2094054.

49. Webb CJ, Zakian VA. Identification and characterization of the Schizosaccharomyces pombe TER1 telomerase RNA. Nat Struct Mol Biol. 2008;15(1):34–42. Epub 2007/12/25. doi: 10.1038/nsmb1354. PubMed PMID: 18157149; PubMed Central PMCID: PMCPMC2703720.

50. Waldl M, Thiel BC, Ochsenreiter R, Holzenleiter A, de Araujo Oliveira JV, Walter M, et al. TERribly Difficult: Searching for Telomerase RNAs in Saccharomycetes. Genes (Basel). 2018;9(8). Epub 2018/07/28. doi: 10.3390/genes9080372. PubMed PMID: 30049970; PubMed Central PMCID: PMCPMC6115765.

51. Egan ED, Collins K. Biogenesis of telomerase ribonucleoproteins. RNA. 2012;18(10):1747–59. Epub 2012/08/10. doi: 10.1261/rna.034629.112. PubMed PMID: 22875809; PubMed Central PMCID: PMCPMC3446700.

52. Qi X, Rand DP, Podlevsky JD, Li Y, Mosig A, Stadler PF, et al. Prevalent and distinct spliceosomal 3’-end processing mechanisms for fungal telomerase RNA. Nat Commun. 2015;6:6105. Epub 2015/01/20. doi: 10.1038/ncomms7105. PubMed PMID: 25598218; PubMed Central PMCID: PMCPMC4299825.

53. Meyne J, Ratliff RL, Moyzis RK. Conservation of the human telomere sequence (TTAGGG)n among vertebrates. Proc Natl Acad Sci U S A. 1989;86(18):7049–53. Epub 1989/09/01. doi: 10.1073/pnas.86.18.7049. PubMed PMID: 2780561; PubMed Central PMCID: PMCPMC297991.

54. Podlevsky JD, Chen JJ. Evolutionary perspectives of telomerase RNA structure and function. RNA Biol. 2016;13(8):720–32. Epub 2016/07/01. doi: 10.1080/15476286.2016.1205768. PubMed PMID: 27359343; PubMed Central PMCID: PMCPMC4993307.

55. Thompson JD, Higgins DG, Gibson TJ. CLUSTAL W: improving the sensitivity of progressive multiple sequence alignment through sequence weighting, position-specific gap penalties and weight matrix choice. Nucleic Acids Res. 1994;22(22):4673–80. Epub 1994/11/11. doi: 10.1093/nar/22.22.4673. PubMed PMID: 7984417; PubMed Central PMCID: PMCPMC308517.

